# Affinity Purification: From Interactome Analysis to Targeted Protein Enrichment

**DOI:** 10.64898/2026.03.02.708958

**Authors:** Amalia Kontochristou, Nika Šimičić, Anouk M. Rijs, Melissa Baerenfaenger

**Affiliations:** Division of Bioanalytical Chemistry, Department of Chemistry and Pharmaceutical Sciences, Amsterdam Institute of Molecular and Life Sciences, Vrije Universiteit Amsterdam, 1081 HV Amsterdam, The Netherlands; Centre for Analytical Sciences Amsterdam (CASA), The Netherlands; Center for Proteomics and Metabolomics, Leiden University Medical Center, 2333 ZA, Leiden, The Netherlands

**Keywords:** Affinity purification, Chaperone, Clusterin, Human plasma, Interactome, Bottom-up proteomics

## Abstract

Extracellular chaperones engage in extensive protein–protein interactions, complicating their selective enrichment and characterisation in complex biological fluids such as human plasma. Consequently, affinity purification workflows are often employed for their analysis. However, the extensive interaction networks of chaperones make purification conditions particularly important, as they can strongly influence the balance between target selectivity and preservation of biologically relevant interactions. In the present study, we systematically evaluated how purification conditions shape affinity purification outcomes for extracellular chaperones using clusterin as a model system. Affinity purification was performed under conditions ranging from interaction-preserving to interaction-disruptive, followed by bottom-up proteomic analysis to assess enrichment and co-purification profiles. Under interaction-preserving conditions, clusterin was enriched together with a broad range of associated proteins, enabling characterisation of its plasma interactome and providing insight into its biological context. Increasing purification stringency progressively reduced co-purification while maintaining clusterin enrichment. Under the most stringent conditions, co-purifying proteins were minimised, enabling more selective isolation of clusterin. Together, these findings demonstrate that affinity purification outcomes can be systematically directed toward either interactome characterisation or selective target enrichment through controlled adjustment of purification conditions. This work establishes a framework for tailoring affinity purification strategies for extracellular chaperones and other highly interactive proteins.

## 1. Introduction

The human proteome is highly complex, comprising thousands of proteins that span a broad dynamic range of abundances [1]. A substantial proportion of the proteome exists within transient and stable protein complexes that regulate biological activity, introducing an additional level of complexity through extensive protein–protein interactions [2]. This complexity is particularly pronounced in biological fluids such as plasma, where a small number of highly abundant proteins, including albumin and immunoglobulin G, account for more than 90% of the total protein content, while many low-abundance proteins are present at concentrations several orders of magnitude lower [3,4]. This dynamic range presents a major challenge for the analysis of low-abundance proteins, as their detection is often hindered by highly abundant species and by their association with interacting partners [5,6]. To enable their analysis, selective enrichment strategies are required, among which affinity purification approaches are widely used for their high binding specificity.

Two main affinity-based strategies are commonly employed for the analysis of low-abundance proteins in human plasma [7]. The first is affinity immunodepletion, in which highly abundant proteins, like albumin, are selectively removed from the sample to increase the relative abundance of low-abundance species. However, this approach has shown limited effectiveness, as low-abundance proteins often form complexes with high-abundance species and may therefore be co-depleted [7]. The second strategy is affinity capture of a specific target protein. This approach has been used for both the selective isolation of a protein of interest and for the characterisation of protein–protein interactions in complex biological systems. In the latter case, a bait protein is isolated along with its binding partners, enabling the detection of stimulus- and condition-dependent protein associations [8–10]. While this is highly beneficial for elucidating protein function, it often leads to extensive co-purification of interacting proteins.

This effect is particularly pronounced for chaperone proteins, which, by definition, bind a broad range of client proteins, thereby increasing the likelihood of extensive co-purification [11]. Chaperones bind to exposed hydrophobic regions of misfolded or stress-damaged proteins and either stabilise them or promote their degradation, thereby maintaining proteostasis [12,13]. By shielding these hydrophobic segments, chaperones suppress protein aggregation and limit the pathological accumulation of proteins in cells and biofluids [14,15]. Clusterin (CLUS) is a glycosylated chaperone protein that is widely distributed across tissues and biofluids, including plasma. As a chaperone, clusterin is involved in various biological processes [16–19], including modulation of inflammation through regulation of the complement system [20], regulation of cell survival and apoptosis-related pathways [19] and lipid transport via high-density lipoprotein (HDL) particles, as well as protein transport across the blood-brain barrier [21–23].

In plasma, where clusterin is present at relatively low abundance, affinity purification is typically employed for protein detection and characterisation, particularly due to its extensive interaction profile [24]. Previous studies have utilised a range of affinity-based strategies, often combined with chromatographic approaches such as size-exclusion, ion-exchange, and hydrophobic interaction chromatography, to enrich for clusterin in biological samples [25–32]. It is noteworthy that these studies frequently reported the co-purification of associated proteins, albeit using affinity purification in combination with subsequent purification steps, which reflects the significant influence of chaperone interactions on the purification outcome [31].

The present study describes how affinity purification conditions can be systematically modulated to direct the analytical outcome toward either preservation of protein interaction networks or more selective target enrichment, thereby providing complementary information on a target protein. Using clusterin as a model extracellular chaperone, we demonstrate that controlled adjustment of affinity purification conditions enables a transition between interaction-preserving and interaction-disruptive regimes. For this purpose, clusterin was enriched from pooled human plasma using a clusterin-specific affinity resin based on single-domain antibody fragments and analysed by bottom-up proteomics. By systematically varying key conditions, the outcome of affinity purification can be directed towards either interactome characterisation or selective protein isolation. These results highlight the versatile nature of affinity purification and the critical role of purification conditions in determining its outcome.

## 2. Materials and Methods

### 2.1 Affinity purification of clusterin

Pooled human plasma (S4180-100) was sourced from Biowest SAS (Nuaillé, France). For the purification of clusterin from 50 μL of human pooled plasma, 100 μL of POROS− CaptureSelect− ClusterinClear Affinity Resin (10 mL), kindly provided by Thermo Fisher Scientific (Carlsbad, CA, USA), was used. The clusterin affinity resin was transferred into a Pierce− Micro-Spin Column supplied by Thermo Fisher Scientific (Rockford, USA). Before applying the plasma sample, the resin was first washed with Milli-Q water and equilibrated with Dulbecco’s Phosphate Buffered Saline (DPBS, pH 7.4) supplied by Thermo Fisher Scientific (Paisley, UK). Milli-Q water was produced by a Millipore purification system (Millipore, Amsterdam, The Netherlands). Plasma was then diluted 1:1 with DPBS and loaded onto the resin. To remove the unbound proteins, the resin was washed multiple times with DPBS buffer and Milli-Q water. Following the washing steps, clusterin was eluted from the affinity material with 100 μL of glycine buffer (pH 2.3), and the eluate was neutralised with 40 μL of Tris(hydroxymethyl)aminomethane (Tris) buffer (pH 9.0), both supplied by Sigma-Aldrich (St. Louis, USA). The sample was then dried under vacuum and stored at −20 °C before bottom-up proteomics analysis. The different purification protocols used are summarised in the Supplementary Material (Table S1).

### 2.2. Bottom-up proteomics

The clusterin-enriched sample was subjected to tryptic digestion. For this, the sample was redissolved in 5 μL of 5 M guanidine hydrochloride and 10 μL of 50 mM DL-dithiothreitol, followed by an incubation at 37 °C with shaking at 650 rpm for 20 minutes. Subsequently, 10 μL of 50 mM iodoacetamide was added, followed by a 60-minute incubation in the dark. Then, 60 μL of 50 mM ammonium bicarbonate and 1 μL of Trypsin/Lys-C (1 mg/mL), supplied by Promega Benelux B.V. (Leiden, The Netherlands), were added, and the sample was incubated overnight at 37 °C with shaking at 650 rpm. Trypsin was inactivated by incubation at 90 °C for five minutes. After tryptic digestion, the sample was stored at −20 °C prior to analysis.

Liquid chromatography-tandem mass spectrometry analysis was performed using an Agilent 1290 Infinity II system (Agilent Technologies Netherlands B.V., Middelburg, The Netherlands) coupled to a trapped ion mobility quadrupole time-of-flight (timsTOF Pro 2) mass spectrometer equipped with an electrospray ionisation (ESI) source operating in positive ion mode (Bruker Daltonics, Bremen, Germany). LC separation was performed at 55 °C with a flow rate of 80 μL/min using solvent A (0.1% formic acid (FA) in Milli-Q water) and solvent B (0.1% FA in acetonitrile). The column used was an Acclaim PepMap 100 C18 reversed-phase column (1 mm ID × 150 mm length, 3 μm particles, 100 Å pore size) supplied by Thermo Fisher Scientific (Ermelo, The Netherlands). The gradient was set as follows: 0 min, 3% B; 2 min, 3% B; 37.5 min, 40% B; 38 min, 90% B; 43.5 min, 90% B; 44 min, 2% B, and 60 min, 2% B. Prior to sample injection, mass and mobility calibration were performed using Agilent Technologies ESI-L tuning mix ions. For MS analysis, the DDA PASEF-standard_1.1sec_cycletime.m Bruker application method was used, with adapted ESI source settings [33]. More information on the MS settings is provided in the Supplementary Material (Table S2).

### 2.3 Data Processing

For data analysis, FragPipe v23.0, MSFragger version 4.3, and Philosopher v5.1.1 were used [34]. Label-free quantification with match between runs (LFQ-MBR) was performed using the IonQuant workflow (version 1.11.11). The UniProt ID UP000005640 database downloaded on 9 December 2023 was used. Precursor mass tolerance was set at ± 20 ppm, and trypsin and Lys-C were selected as the enzymes for proteolytic digestion. Carbamidomethylation of cysteine was set as a fixed modification, while methionine oxidation, pyroglutamic acid formation, loss of ammonia at the peptide N-terminus, and N-terminal acetylation were specified as variable modifications. For MS1 quantification, the MaxLFQ min ions were set to 3, the MBR ion FDR was set to 0.01, and peptide-protein uniqueness was restricted to unique only.

The generated data was processed and visualised using in-house developed Python scripts, which are available upon request. To visualise the distribution of protein abundances across the samples (Protein Abundance Plots), MaxLFQ Intensity values were used. For each protein, the mean MaxLFQ Intensity was calculated across replicates, and proteins with non-positive mean values were excluded. Then, the mean intensities were log_10_-transformed, sorted in descending order, and plotted as abundance rank versus log_10_ (MaxLFQ Intensity) using matplotlib.pyplot library version 3.10.0.

For the volcano plots, the MaxLFQ Intensity of proteins was log_2_-transformed, and median-centred normalisation was applied per condition to correct for systematic variation. Missing values were imputed per condition using a normal distribution with a downshift of 1.8 standard deviations and a width of 0.3 times the standard deviation of the observed values, following a Perseus-style approach. Welch’s t-test was used to compare conditions, and log_2_ fold changes (FC) were calculated from the mean normalised intensities. Volcano plots were generated using the seaborn and matplotlib libraries, with log_2_FC on the x-axis and –log_10_ (p-value) on the y-axis. Proteins with |log_2_FC| ≥ 1 and p < 0.05 were considered significantly enriched.

To define the interactome of clusterin, proteins significantly enriched after clusterin purification relative to non-enriched human plasma, as identified by volcano plot analysis, were selected and analysed using the STRING database version 12.0. Details of the STRING parameters and settings used for network generation are provided in the Supplementary Material (Table S3).

For bar plot generation, the relative abundance of clusterin was calculated for each replicate as the percentage of its MaxLFQ Intensity relative to the total MaxLFQ Intensity across all proteins within the sample. Relative clusterin abundances were summarised across experimental conditions and visualised as bar plots using seaborn version 0.13.2 and matplotlib.pyplot libraries.

Lastly, to visualise global protein enrichment, mean MaxLFQ Intensities were calculated per protein across replicates for each condition. Log_2_FC were then determined by comparing enriched samples (Protocol 1–3) to the non-enriched plasma sample using a pseudocount. The distribution of log_2_FC values was visualised using boxplots overlaid with jittered points representing individual proteins.

## 3. Results and Discussion

### 3.1 Affinity Purification Workflow

The affinity purification workflow was implemented using a clusterin-specific affinity resin in a solid-phase extraction (SPE) format. The workflow consisted of three main steps: capture, washing and elution (Figure 1). Following sample loading, clusterin was retained on the resin through antibody-antigen interactions, while non-bound proteins were removed during the washing step. Elution was achieved by lowering the pH to disrupt antibody-antigen interactions.

**Figure 1.**
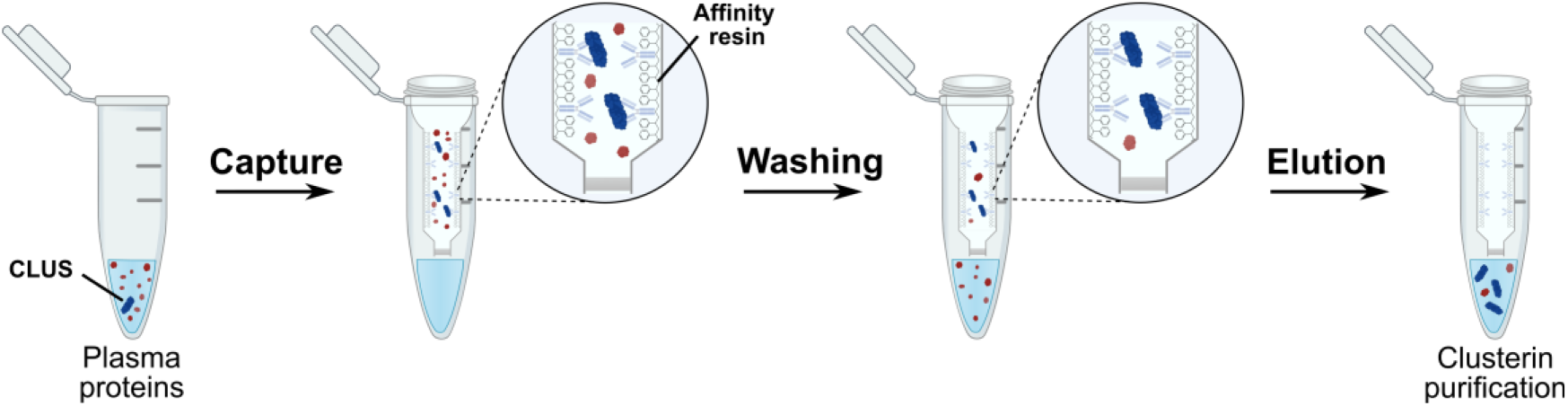
Clusterin purification workflow using POROS− CaptureSelect− ClusterinClear affinity resin. The purification was performed in a spin column and involved three sequential steps. Capture: Human plasma was loaded onto the affinity resin, where clusterin (CLUS) bound selectively to the immobilised antibody. Washing: Unbound plasma proteins (in red) were removed. Elution: The interaction between clusterin and the antibody was disrupted, resulting in protein elution.

To evaluate the effect of purification conditions on clusterin enrichment, affinity purification was performed under a range of conditions varying from interaction-preserving to interaction-disruptive. Enriched samples were analysed using label-free quantification bottom-up proteomics, enabling assessment of both clusterin recovery and the extent of co-purification.

#### 3.2. Interactome Analysis

The initial affinity purification protocol evaluated in this study (Protocol 1) was established according to the supplier’s instructions for the clusterin affinity resin adapted for an SPE approach. Similar conditions have been applied in previous studies for the purification of clusterin-containing complexes [35], as well as for the purification of other human proteins [36,37]. Protocol 1 employed near-physiological conditions during equilibration and washing (DPBS, pH 7.0–7.5), thereby favouring preservation of protein–protein interactions. Elution was induced using a low-pH solution (0.1 M glycine, pH 2.3), disrupting the clusterin-antibody interaction and releasing the target protein together with associated proteins.

To assess the outcome of the purification, the relative abundance of proteins in the enriched sample was compared with that in non-enriched plasma. For this purpose, log10-transformed MaxLFQ Intensities of all identified proteins were plotted against abundance rank to visualise the dynamic range of proteins in the sample. As shown in Figure 2, Protocol 1 resulted in substantial clusterin enrichment relative to the non-enriched plasma, reflected by a pronounced shift in its abundance rank from a mid-abundance position (rank ∼45–50) to among the top-ranked proteins (rank <10). At the same time, a reduction in the total number of detected proteins was observed, reflecting partial removal of background plasma proteins while retaining a subset of co-purified species.

**Figure 2.**
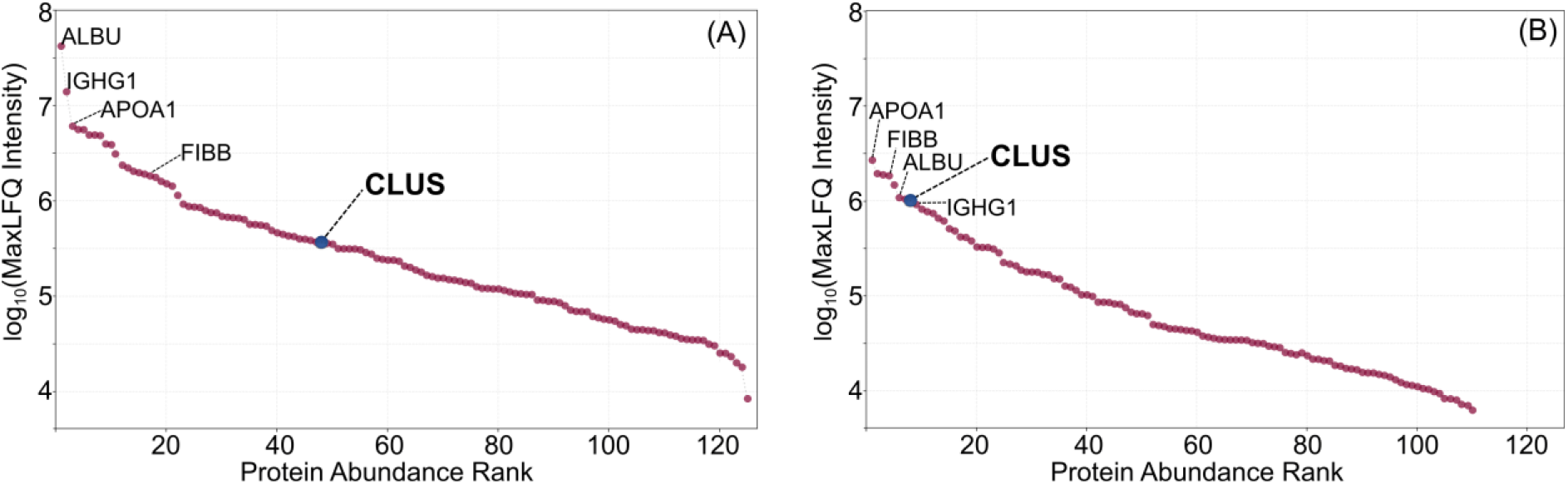
Protein abundance profiles of (A) non-enriched and (B) clusterin-enriched human plasma following Protocol 1. Proteins were ranked by abundance on the x-axis, while the y-axis displays the average log_10_ (MaxLFQ Intensity) for each protein. Each point represents an individual protein. Selected highly abundant proteins (ALBU, IGHG1, APOA1, and FIBB) are highlighted with clusterin (CLUS) shown in blue.

Despite successful clusterin enrichment, a substantial number of proteins co-purified with clusterin, and the dynamic range of protein abundances was only modestly reduced following enrichment (Figure 2B). Consistently, several highly abundant plasma proteins, including albumin (ALBU), immunoglobulin G (IGHG1), apolipoprotein A1 (APOA1), and fibrinogen beta chain (FIBB), remained among the most abundant species after purification.

To identify proteins associated with clusterin under these purification conditions, the relative abundance of proteins in the enriched sample was compared to that in non-enriched plasma using a volcano plot approach (Figure 3A). It was assessed based on both fold change and statistical significance to discriminate between background highly abundant plasma proteins and proteins co-purified with clusterin. Proteins showing a positive log2 fold change (enriched vs non-enriched) and meeting defined significance threshold (p-value < 0.05) were considered candidate clusterin-associated proteins. The resulting protein set was subsequently analysed using the STRING network analysis [38] to visualise predicted and experimentally validated interactions between clusterin and other plasma proteins, based on both physical and functional associations (Figure 3B).

**Figure 3.**
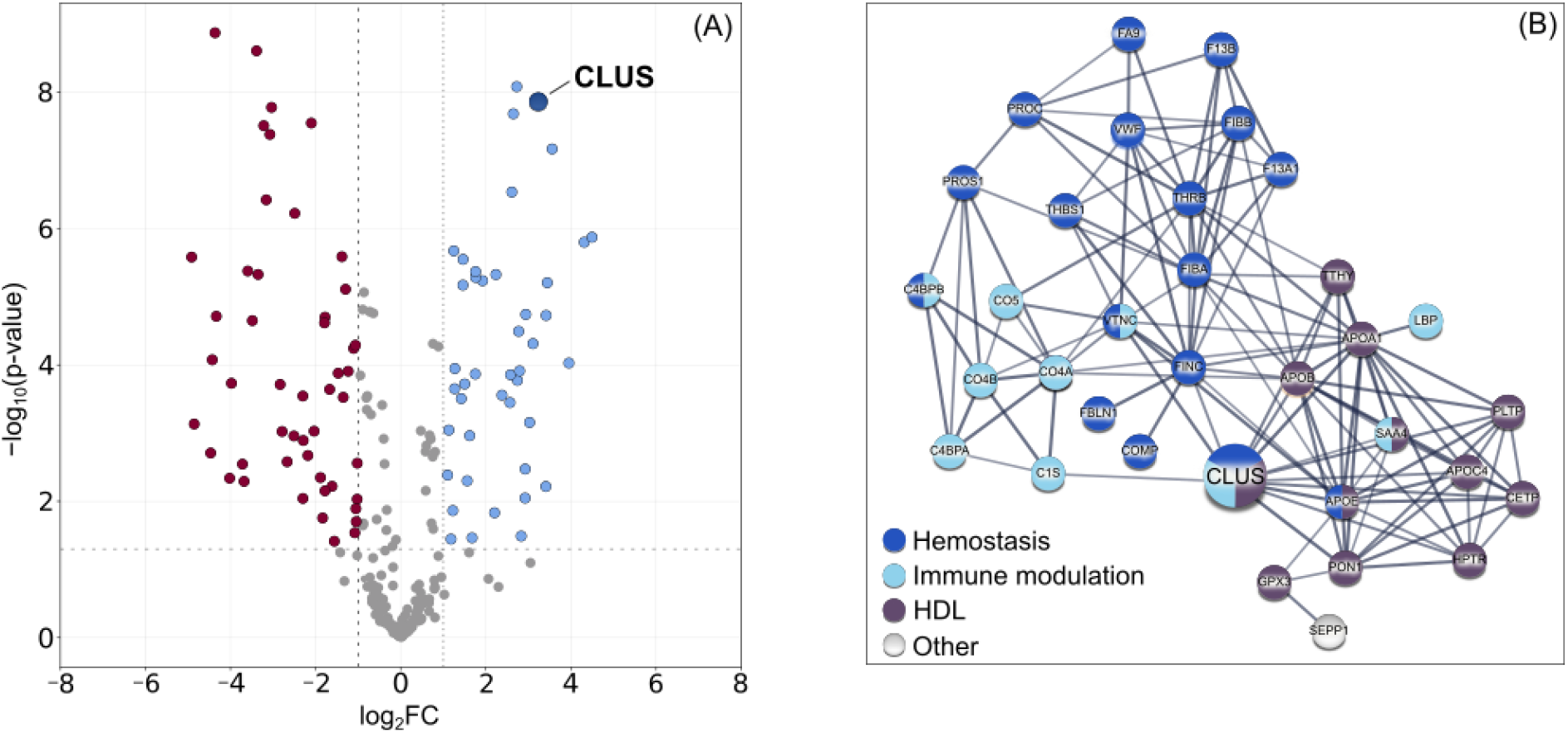
(A) Volcano plot comparing non-enriched and enriched human plasma. Log2FC (x-axis) is plotted against –log_10_ (p-value) (y-axis). Proteins significantly enriched following affinity purification (Protocol 1) are highlighted in blue, whereas proteins more abundant in non-enriched plasma are shown in red. (B) Clusterin interactome in human plasma, visualised using the STRING network database. The network comprises 40 proteins significantly enriched, following clusterin purification using Protocol 1, and is based on known and predicted protein–protein interactions. Edges represent confidence scores between proteins, with a minimum required interaction score of 0.700 (high confidence), and edge thickness corresponds to interaction strength. Protein nodes were grouped according to biological function: immune modulation (light blue), HDL particles (brawn), and hemostasis pathway (dark blue). SEPP1 (white) was the only protein not assigned to the aforementioned groups.

The STRING network analysis revealed that clusterin is part of an extensive protein–protein interaction network, comprising both direct and indirect associations. The identified proteins could be broadly grouped into three categories based on their biological roles: hemostasis-related proteins, immune modulation-associated proteins, and components of high-density lipoprotein (HDL) particles. HDL particles are complexes composed mainly of phospholipids, cholesterol, and apolipoproteins, whose primary function is lipid transport between cells [39]. Clusterin, also known as apolipoprotein J, is a known component of HDL and has been reported to associate with lipoprotein particles [40]. Consistent with this, the identified interactome comprised several HDL-related proteins [35,40–47].

In addition to HDL-associated proteins, numerous proteins related to hemostasis were also identified within the clusterin interactome [48–51]. Although clusterin is not a direct component of the coagulation cascade, extracellular chaperones have been reported to associate with fibrin clots, where they contribute to the stabilisation of unfolded or misfolded proteins, thereby delaying or inhibiting clot formation and exerting a potential anticoagulant effect [51–53].

Proteins involved in immune response pathways also represented a prominent component of the identified interactome, comprising acute-phase proteins such as SAA4 and LBP [54], as well as components of the complement cascade [55]. The complement system is part of innate immunity, responsible for tagging pathogens for destruction, activating immune cells, and directly eliminating pathogens through the membrane attack complex (MAC) mechanism [56,57]. Clusterin has been reported to inhibit MAC formation, thereby protecting host cells from complement-mediated damage and preventing the lysis of healthy, non-target cells, together with another protein present in the interactome, vitronectin (VTNC) [58].

Notably, most proteins identified within the interactome have previously been reported to associate with clusterin, supporting the biological relevance of the observed interactions [38–48]. This consistency suggests that the detected proteins predominantly reflect established interactions with clusterin or clusterin-containing complexes rather than non-selective enrichment of abundant plasma components. It is also important to note that, although the identified proteins could be broadly grouped into three functional categories, these classifications were not mutually exclusive. Interactions between proteins across different groups were also observed, reflecting the complexity of protein–protein interaction networks, where proteins and multiprotein complexes often participate in multiple interconnected biological processes. Such overlap is also consistent with the diverse biological functions previously associated with clusterin [38–48]. This complexity may explain the presence of SEPP1 within the interactome, despite not clearly belonging to any of the three major groups. SEPP1 is functionally connected to GPX3, as both proteins are known to be involved in selenium-dependent redox homeostasis [59]. The complete list of proteins in the network is provided in the Supplementary Material (Table S4).

These findings indicate that the applied purification conditions favour preservation of clusterin-associated protein complexes, enabling capture of its interaction network in a near-physiological state. However, the extent of co-purification observed under these conditions highlights the need to tune purification conditions to shift the workflow from interactome characterisation towards more selective target protein isolation.

#### 3.3. Protein Isolation

To transition from interactome characterisation towards more selective protein isolation, the affinity purification conditions were systematically adjusted to minimise co-purification while maintaining efficient clusterin enrichment. Individual workflow parameters were varied independently to assess their specific contribution to the purification outcome.

As an initial step, the effect of incubation time during the capture stage was evaluated by varying the interaction time between the plasma sample and the clusterin affinity resin under controlled conditions (650 rpm, 4 °C). Incubation times ranging from no incubation to 2 hours (0, 30 min, 1 h, 2 h) were tested to determine whether prolonged interaction enhanced clusterin binding. However, no significant differences in clusterin enrichment were observed across the tested conditions (Figure S1), indicating that efficient binding occurs within a very short period of time.

Following optimisation of the capture step conditions, the composition of the washing buffer was systematically varied to reduce co-purification and improve clusterin isolation. Specifically, the concentrations of NaCl and Tween 20 in the DPBS washing buffer were adjusted to modulate ionic and hydrophobic protein–protein interactions.

NaCl was first evaluated for its ability to suppress ionic interactions through charge shielding of protein surfaces. NaCl is a native component of DPBS and dissociates into Na+ and Cl− ions, both of which are positioned near the centre of the Hofmeister Series and are therefore expected to exert minimal effects on protein structure and stability [60,61]. Importantly, all purification conditions evaluated were selected to minimise disruption of protein conformation and preserve binding to the immobilised antibody of the affinity resin.

Subsequently, Tween 20 was assessed as an additional washing buffer additive to target hydrophobic interactions. As a non-ionic detergent, Tween 20 can associate with exposed hydrophobic regions of proteins, thereby reducing aggregation and limiting non-specific protein–protein interactions involving clusterin [62]. Also, Tween 20 has been reported to exert a “renaturating effect” on proteins, referring to its capacity to maintain proteins in solution without inducing denaturation [63].

When evaluated individually, Tween 20 resulted in a more pronounced reduction in co-purified proteins compared to NaCl alone. This observation is consistent with the chaperone function of clusterin, which mainly involves interactions with exposed hydrophobic regions of client proteins, suggesting that hydrophobic interactions play a major role in the observed co-purification. However, the combined use of both NaCl and Tween 20 yielded the highest degree of purification (Figure S2), indicating that both ionic and hydrophobic interactions contribute to the observed co-purification and must be simultaneously disrupted to achieve more selective isolation.

Multiple combinations of NaCl and Tween 20 concentrations were evaluated by varying one component while keeping the other constant, to identify conditions that maximised clusterin enrichment. As shown in Figure 4A, the relative abundance of clusterin increased with increasing NaCl concentration, reaching a maximum at 0.75 M NaCl. Further increases to 1.0 M and 1.5 M NaCl did not improve purification efficiency and instead resulted in a decline in clusterin relative abundance. This trend is consistent with the salting-in effect, which describes how a moderate increase in ionic strength can shield protein surface charges, reducing protein–protein ionic interactions. Although at higher salt concentrations, the competition between salt ions and proteins for water molecules can reduce protein solubility by limiting hydration shell formation around proteins, thereby promoting protein–protein interactions and precipitation [64]. Based on these observations, higher NaCl concentrations were not tested to avoid potential protein precipitation effects and to maintain compatibility with downstream mass spectrometry analysis.

**Figure 4.**
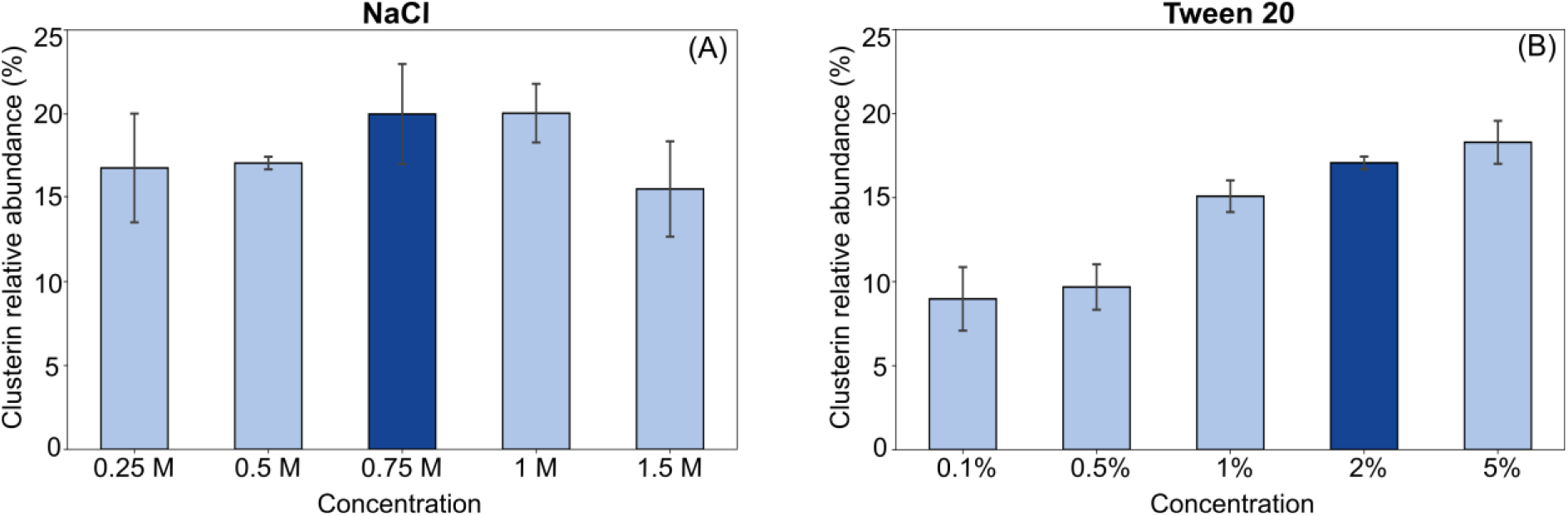
Optimisation of washing buffer composition for clusterin purification. Bar plots show the effect of varying concentrations of (A) NaCl and (B) Tween 20 in the washing buffer on the relative abundance of clusterin. The x-axes indicate the concentrations of NaCl and Tween 20 tested. The y-axes represent the ratio of the log_10_ (MaxLFQ Intensity) of clusterin to the log_10_ (MaxLFQ Intensity) of all proteins in the sample, reflecting relative clusterin abundance. During Tween 20 optimisation, the NaCl concentration was held constant at 0.5 M, while during NaCl optimisation, the Tween 20 concentration was maintained at 2%. Error bars represent standard deviation across three replicates.

In the case of Tween 20 (Figure 4B), increasing its concentration while keeping the NaCl level constant resulted in a progressive improvement in clusterin purification. This effect is consistent with the disruption of hydrophobic protein–protein interactions at higher detergent levels [65]. The most substantial improvement was observed between 0.5% and 2% Tween 20. Specifically, increasing the concentration from 0.5% to 1% resulted in a 1.6-fold increase in clusterin relative abundance, followed by a further 1.1-fold increase at 2%. Increasing the concentration to 5% yielded only a minor additional improvement. Considering the limited gain at higher detergent concentrations, and to avoid potential protein denaturation [66], 2% Tween 20 was selected as the optimal concentration.

Based on these findings, the purification protocol was refined as Protocol 2, incorporating a total of eight washing steps. Following sample loading, the first two washes were performed with DPBS supplemented with 0.75 M NaCl and 2% Tween 20 to minimise ionic and hydrophobic interactions. This was followed by two washes with DPBS containing 0.75 M NaCl only, to further reduce protein co-purification. Subsequently, four consecutive washes with Milli-Q water were implemented to remove remaining Tween 20 and NaCl, ensuring MS compatibility and facilitating the transition to low-pH elution conditions.

Following optimisation of the washing step, the elution conditions were also modified to test the effect on the purification outcome. While elution in Protocols 1 and 2 was performed using a glycine buffer (pH 2.3), Protocol 3 incorporated the addition of 0.5 M NaCl to the glycine solution (Protocol 3). The inclusion of NaCl resulted in a significant decrease in the number of co-purified plasma proteins (Figure S2D). This effect is consistent with changes in protein solubility under acidic and high-ionic-strength conditions. At low pH, many plasma proteins carry a net positive charge, leading to electrostatic repulsion that helps maintain solubility [67]. The addition of salt can shield these surface charges, reduce electrostatic repulsion and promote protein–protein interactions, which may lead to aggregation or precipitation and consequently lower protein solubility [64].

Comparison of Protocols 1-3 revealed clear differences in both clusterin enrichment and the extent of protein co-purification, as illustrated in Figure 5. Using interaction-preserving conditions (Protocol 1), affinity purification resulted in extensive co-enrichment of a large number of proteins with clusterin, making this approach well-suited for monitoring the clusterin interactome in complex biological samples.

**Figure 5.**
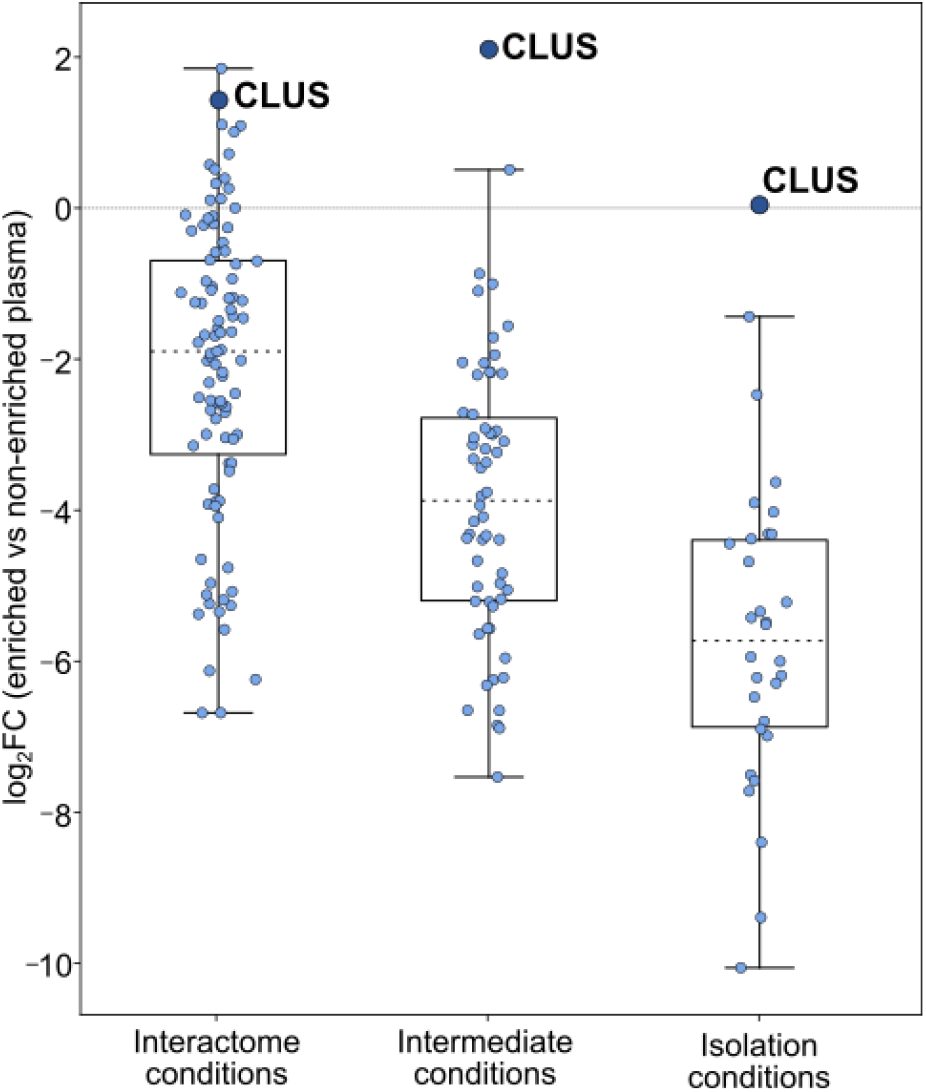
Global shift in protein enrichment across purification conditions. Log_2_ fold change (enriched vs non-enriched plasma) is shown for proteins identified under three purification protocols: interaction-preserving conditions (Protocol 1), intermediate conditions (Protocol 2), and isolation conditions (Protocol 3). Each point represents an individual protein, and boxplots indicate the distribution of enrichment values. The number of proteins decreases progressively across conditions, reflecting reduced co-purification. Clusterin (CLUS) is highlighted. The shift towards lower fold change values from Protocol 1 to Protocol 3 indicates increased purification selectivity.

Introduction of intermediate, more stringent conditions designed to minimise protein–protein interactions (Protocol 2) led to a substantial decrease in co-purification. This was accompanied by an overall shift towards lower log_2_ fold change values (Figure 5), indicating a reduction in the relative abundance of co-purified proteins in the enriched sample compared to non-enriched plasma and suggesting improved disruption of non-specific interactions, while clusterin enrichment increased under these conditions. Apart from clusterin, paraoxonase 1 (PON1) was the only protein that remained more abundant in the enriched sample compared to non-enriched plasma, consistent with its reported association with clusterin [68]. Application of even more stringent purification conditions (Protocol 3) resulted in a further reduction in co-purified proteins, approximately by half. Notably, clusterin remained enriched under all conditions, although its relative enrichment decreases under the most stringent conditions, highlighting a trade-off between purification selectivity and targeted protein recovery.

## 4. Conclusion

Affinity purification is a versatile technique that can provide multiple layers of information on proteins of interest. Its outcome is not fixed, but strongly dependent on the applied purification conditions, which determine whether protein–protein interactions are preserved or disrupted. By systematically modulating these conditions throughout the workflow, different types of information can be obtained. For highly interactive proteins such as clusterin, which participate in extensive protein– protein interaction networks within complex biological mixtures, affinity purification in combination with bottom-up proteomics is particularly valuable, as it enables both functional insights through interactome analysis and selective target isolation. By tuning interaction-preserving and interaction-disruptive parameters, the workflow can be directed towards either interactome mapping or targeted enrichment of the protein of interest. This approach improves access to clusterin in complex biofluids and enables detailed analysis of clusterin-associated proteins in human plasma. Overall, this work highlights the importance of purification design and demonstrates how affinity purification workflows can be rationally tailored to achieve different analytical outcomes. While demonstrated here for clusterin, this approach provides a generalised framework for designing protein purification strategies in complex biological systems.

## Supporting information

All combined_protein files generated by FragPipe search for all conditions tested.

## CRediT AUTHORSHIP CONTRIBUTION STATEMENT

The manuscript was written with contributions from all authors. All authors have approved the final version. Amalia Kontochristou contributed to the conceptualisation, data curation, formal analysis, investigation, methodology, validation, visualisation and writing - original draft. Nika Šimičić contributed to conceptualisation and methodology. Anouk M. Rijs contributed to funding acquisition, supervision, and writing - review and editing. Melissa Baerenfaenger contributed to conceptualisation, funding acquisition, methodology, project administration, supervision, visualisation, and writing - review and editing.

## ACKNOWLEDGEMENTS

The authors thank Nicolas Laroudie and Dr. Maria Bartel from Thermo Fisher Scientific for their support and input on the development of the clusterin purification workflows. We would also like to thank Dr. Julien Slagboom for his helpful suggestions on developing the affinity purification protocol, as well as Raya Sadighi and Chuck van der Veen for maintaining the LC and MS instruments used in this study.

## DATA AVAILABILITY STATEMENT

The mass spectrometry proteomics data and the FragPipe workflow used have been deposited to the ProteomeXchange Consortium via the PRIDE partner repository with the dataset identifier PXD075110.

## SUPPLEMENTARY MATERIAL

The supplementary material includes: Detailed experimental procedures, including purification workflow conditions (Table S1); mass spectrometry method parameters (Table S2); STRING network analysis parameters (Table S3); and a complete list of proteins identified in the clusterin interactome (Table S4); protein abundance plots comparing post sample loading incubation times (Figure S1); protein abundance plots following optimisation of the conditions of the washing and elution steps (Figure S2) (PDF). All combined_protein.tsv output files generated by FragPipe for each experimental condition tested (XLSX).

## ABBREVIATIONS

CLUS: Clusterin
DDA-PASEF: Data Dependent - Parallel Accumulation Serial Fragmentation
DPBS: Dulbecco’s Phosphate Buffered Saline
ESI: Electrospray Ionisation
FA: Formic acid
FC: Fold Change
FDR: False Discovery Rate
HDL: High-density lipoprotein
HPLC: High Performance Liquid Chromatography
LC-MS/MS: Liquid Chromatography Tandem Mass Spectrometry
LFQ-MBR: Label Free Quantification - Match Between Runs
MAC: Membrane Attack Complex
MaxLFQ: Maximum Limit of Quantification
SPE: Solid Phase Extraction
STRING: Search Tool for the Retrieval of Interacting Genes/Proteins
timsTOF: Trapped Ion Mobility Time of Flight
ULC: Ultra Performance Liquid.

