## Supplementary material for "Affinity Purification: From Interactome Analysis to Targeted Protein Enrichment": Supplemetary Material

**Table of Contents:**

**Table S1:** Clusterin purification workflows.

**Table S2:** MS method settings.

**Table S3:** STRING network analysis parameters.

**Tables S4:** Summary of proteins identified in clusterin interactome network.

**Figure S1:** Protein abundance plots comparing post-loading incubation times.

**Figure S2:** Protein abundance plots comparing buffer compositions.

**Table S1:** Summary of clusterin purification workflows tested. For each protocol, only NaCl and Tween 20 concentrations are reported, as all other buffer components remained unchanged. The NaCl concentration in DPBS (0.137 M) was adjusted in each protocol to achieve the final NaCl concentrations listed in the table.

| **Protocols** | | **Washing** | **Elution** |
|---|---|---|---|
| 1 | | DPBS *(x3)* | Glycine |
| **2.1** | *a)* | DPBS: 0.5 M NaCl + 0.1% Tween 20 *(x2)* | Glycine |
| | *b)* | DPBS: 0.5 M NaCl *(x2)* | |
| | *c)* | Milli-Q *(x4)* | |
| **2.2** | *a)* | DPBS: 0.5 M NaCl + 0.5% Tween 20 *(x2)* | Glycine |
| | *b)* | DPBS: 0.5 M NaCl *(x2)* | |
| | *c)* | Milli-Q *(x4)* | |
| **2.3** | *a)* | DPBS: 0.5 M NaCl + 1% Tween 20 *(x2)* | Glycine |
| | *b)* | DPBS: 0.5 M NaCl *(x2)* | |
| | *c)* | Milli-Q *(x4)* | |
| **2.4** | *a)* | DPBS: 0.5 M NaCl + 2% Tween 20 *(x2)* | Glycine |
| | *b)* | DPBS: 0.5 M NaCl *(x2)* | |
| | *c)* | Milli-Q *(x4)* | |
| **2.5** | *a)* | DPBS: 0.5 M NaCl + 5% Tween 20 *(x2)* | Glycine |
| | *b)* | DPBS: 0.5 M NaCl *(x2)* | |
| | *c)* | Milli-Q *(x4)* | |
| **2.6** | *a)* | DPBS: 0.25 M NaCl + 2% Tween 20 *(x2)* | Glycine |
| | *b)* | DPBS: 0.25 M NaCl *(x2)* | |
| | *c)* | Milli-Q *(x4)* | |
| **2.7** | *a)* | DPBS: 0.5 M NaCl + 2% Tween 20 *(x2)* | Glycine |
| | *b)* | DPBS: 0.5 M NaCl *(x2)* | |
| | *c)* | Milli-Q *(x4)* | |
| **2** | *a)* | DPBS: 0.75 M NaCl + 2% Tween 20 *(x2)* | Glycine |
| | *b)* | DPBS: 0.75 M NaCl *(x2)* | |
| | *c)* | Milli-Q *(x4)* | |

| | | |
|---|---|---|
| **2.8** | *a)* DPBS: 1 M NaCl + 2% Tween 20 *(x2)* | Glycine |
| | *b)* DPBS: 1 M NaCl *(x2)* | |
| | *c)* Milli-Q *(x4)* | |
| **2.9** | *a)* DPBS: 1.5 M NaCl + 2% Tween 20 *(x2)* | Glycine |
| | *b)* DPBS: 1.5 M NaCl *(x2)* | |
| | *c)* Milli-Q *(x4)* | |
| **2.10** | a) DPBS: 0.137 M NaCl+ 2% Tween 20 *(x2)* | Glycine |
| | b) DPBS: 0.137 M NaCl *(x2)* | |
| | c) Milli-Q *(x4)* | |
| **2.11** | a) DPBS: 0.75 M NaCl *(x2)* | Glycine |
| | b) DPBS: 0.137 M NaCl *(x2)* | |
| | c) Milli-Q *(x4)* | |
| **3** | *a)* DPBS: 0.75 M NaCl + 2% Tween 20 *(x2)* | Glycine + 0.5 M NaCl |
| | *b)* DPBS: 0.75 M NaCl *(x2)* | |
| | *c)* Milli-Q *(x4)* | |

**Table S2:** Setting of the DDA PASEF-standard_1.1sec_cycletime.m MS method.

| **Parameters** | **Values** |
|---|---|
| Capillary voltage [V] | 4500 |
| Nebulizer [Bar] | 0.4 |
| Dry Gas [l/min] | 3.0 |
| Dry Temp [ºC] | 180 |
| Number of PASEF ramps | 10 |
| Total Cycle Time [s] | 1.17 |
| Ramp Time [ms] | 100 |
| Accumulation Time [ms] | 100 |

**Table S3:** String network parameters and network statistics used for STRING analysis.

| **Parameters** | **Values** |
|---|---|
| Number of nodes | 40 |
| Number of edges | 128 |
| Average node degree | 6.4 |
| Average local clustering coefficient | 0.643 |
| Expected number of edges | 4 |
| PPI enrichment p-value | $< 1.0\ e^{-16}$ |

**Table S4:** List of plasma proteins involved in the interactome of clusterin

| **UniProt ID** | **Entry Name** | **Gene Name** | **Description** |
|---|---|---|---|
| P02751 | FINC_HUMAN | FN1 | Fibronectin |
| P01031 | CO5_HUMAN | C5 | Complement C5 |
| P04275 | VWF_HUMAN | VWF | von Willebrand factor |
| P20851 | C4BPB_HUMAN | C4BPB | C4b-binding protein beta chain |
| P04003 | C4BPA_HUMAN | C4BPA | C4b-binding protein alpha chain |
| P02671 | FIBA_HUMAN | FGA | Fibrinogen alpha chain |
| P00488 | F13A_HUMAN | F13A1 | Coagulation factor XIII A chain |
| P10909 | CLUS_HUMAN | CLU | Clusterin |
| P02679 | FIBG_HUMAN | FGG | Fibrinogen gamma chain |
| Q92496 | FHR4_HUMAN | CFHR4 | Complement factor H-related protein 4 |
| P07996 | TSP1_HUMAN | THBS1 | Thrombospondin-1 |
| P55058 | PLTP_HUMAN | PLTP | Phospholipid transfer protein |
| P18428 | LBP_HUMAN | LBP | Lipopolysaccharide-binding protein |
| P02654 | APOC1_HUMAN | APOC1 | Apolipoprotein C-I |
| P02675 | FIBB_HUMAN | FGB | Fibrinogen beta chain |
| P04114 | APOB_HUMAN | APOB | Apolipoprotein B-100 |
| P02743 | PZP_HUMAN | PZP | Pregnancy zone protein |
| P27169 | PON1_HUMAN | PON1 | Serum paraoxonase/arylesterase 1 |

| | | | |
|---|---|---|---|
| P07225 | PROS_HUMAN | PROS1 | Vitamin K-dependent protein S |
| Q9HCB6 | FCGBP_HUMAN | FCGBP | IgGFc-binding protein |
| P07204 | THRB_HUMAN | F2 | Prothrombin |
| P0C0L4 | CO4A_HUMAN | C4A | Complement C4-A |
| P15169 | CBPN_HUMAN | CPB2 | Carboxypeptidase N catalytic chain |
| P01871 | IGHM_HUMAN | IGHM | Immunoglobulin heavy constant mu |
| Q9UHK0 | PHLD_HUMAN | PLEKHD1 | Pleckstrin homology-like domain family A member 1 |
| O43866 | CD5L_HUMAN | CD5L | CD5 antigen-like |
| P22792 | CPN2_HUMAN | CPN2 | Carboxypeptidase N subunit 2 |
| P02766 | TTHY_HUMAN | TTR | Transthyretin |
| P49908 | SEPP1_HUMAN | SELENOP | Selenoprotein P |
| P35542 | SAA4_HUMAN | SAA4 | Serum amyloid A-4 protein |
| P04070 | PROC_HUMAN | PROC | Vitamin K-dependent protein C |
| P01591 | IGJ_HUMAN | IGJ | Immunoglobulin J chain |
| P02647 | APOA1_HUMAN | APOA1 | Apolipoprotein A-I |
| P02649 | APOE_HUMAN | APOE | Apolipoprotein E |
| P0C0L5 | CO4B_HUMAN | C4B | Complement C4-B |
| P06727 | APOA4_HUMAN | APOA4 | Apolipoprotein A-IV |
| P23142 | FBLN1_HUMAN | FBLN1 | Fibulin-1 |
| P04004 | VTNC_HUMAN | VTN | Vitronectin |
| P12259 | FA5_HUMAN | F5 | Coagulation factor V |
| P11597 | CETP_HUMAN | CETP | Cholesteryl ester transfer protein |
| P22352 | GPX3_HUMAN | GPX3 | Glutathione peroxidase 3 |

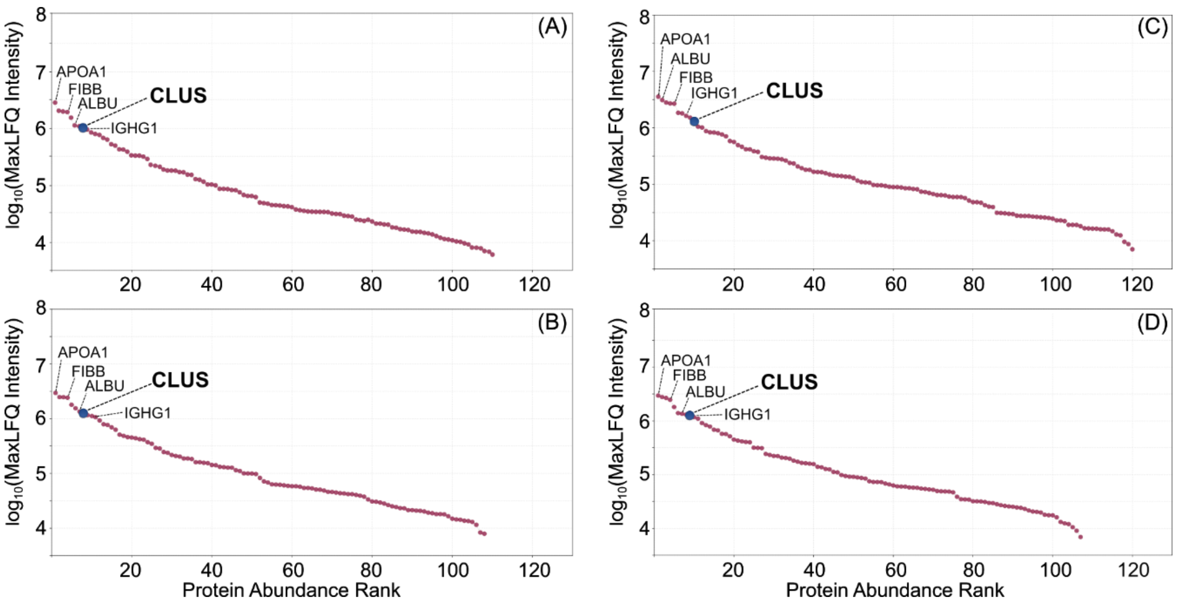


**Figure S1:** Protein abundance plots of clusterin-enriched obtained using Protocol 1, in which the post-loading incubation time was the only parameter varied. Incubation durations after sample loading were: (A) No incubation, (B) 30 minutes, (C) 1 hour, and (D) 2 hours. Proteins are ranked by abundance on the x-axis, while the y-axis displays the average $\log_{10}$(MaxLFQ Intensity) for each protein. Each red dot represents an individual protein, with clusterin (CLUS) highlighted in blue. Increasing the incubation time after sample loading did not substantially improve clusterin enrichment or reduce co-purification.

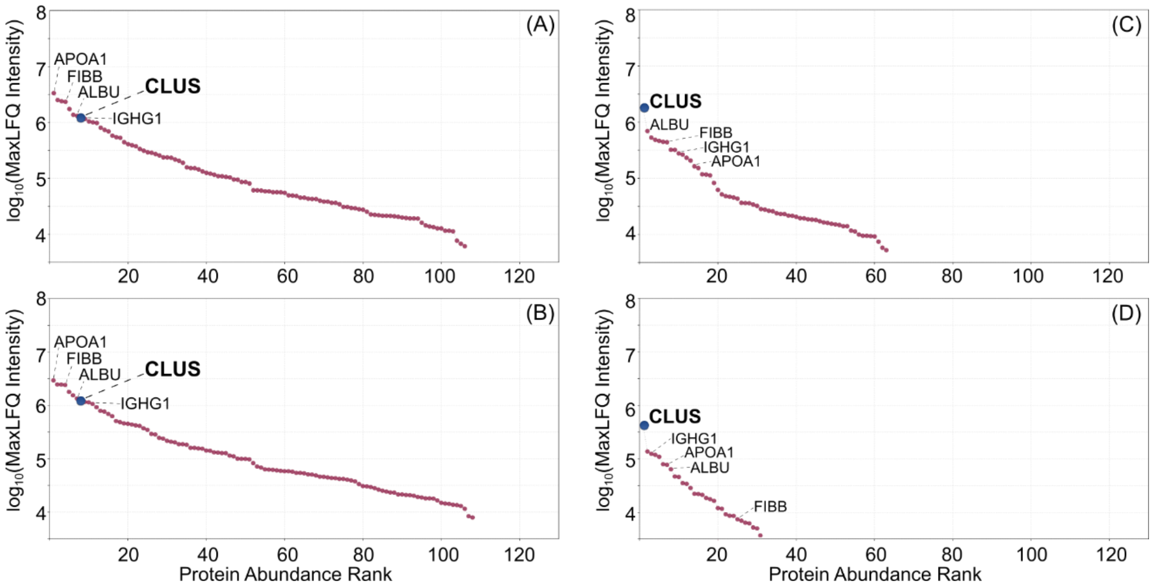


**Figure S2:** Protein abundance in clusterin-enriched samples following optimisation of washing and elution buffer composition. Proteins are ranked by abundance on the x-axis, while the y-axis displays the average

$log_{10}$(MaxLFQ Intensity) for each protein. Each red dot represents an individual protein, with clusterin (CLUS) highlighted in blue. (A) Effect of increasing NaCl concentration in the washing buffer to a final concentration of 0.75 M NaCl (Protocol 2.11). (B) Effect of adding 2% Tween 20 to the washing buffer without additional NaCl (Protocol 2.10). In both conditions, clusterin was enriched but co-purified with several plasma proteins; however, Tween 20 exerted a more pronounced effect than NaCl alone, as reflected by the higher abundance rank of clusterin in panel B. (C) Clusterin enrichment following Protocol 2. (D) Clusterin enrichment following Protocol 3, in which NaCl was added to the elution buffer. While clusterin remained enriched in both protocols, the addition of NaCl to the elution buffer (Protocol 3) substantially reduced co-purification, indicating improved purification efficiency.
